# Efficient Generation of Functional TCRαβ^+^ Cytotoxic T Cells from hiPSCs via Small-Molecule Modulation

**DOI:** 10.64898/2026.03.31.715684

**Authors:** Caroline Kubaczka, Netra Kambli, Roland Windisch, Kelly Yu, Yunliang Zhao, Sharon Wu, Katie Frenis, Morgan T. Walcheck, Marcelo Falchetti, Mohamad Najia, Zachary C LeBlanc, Trista E. North, R Grant Rowe, George Q. Daley, Thorsten M. Schlaeger

**Affiliations:** Stem Cell and Regenerative Biology Program, Boston Children’s Hospital, Boston, MA 02115, USA; Department of Biological Chemistry and Molecular Pharmacology, Harvard Medical School, Boston, MA 02115, USA; Department of Hematology-Oncology, Boston Children’s Hospital, Boston, MA, USA

**Keywords:** pluripotent stem cells, Aryl hydrocarbon receptor (AHR), WNT, GSK, DOT1L, lymphoid development, T cell differentiation, cytotoxic T cells, hematopoiesis, chemical screen, CAR-T cells, cancer immunotherapy, autoimmune disease

## Abstract

Genetically engineered human induced pluripotent stem cells (hiPSCs) represent a promising platform for regenerative medicine and next-generation immunotherapies. While recent advances enable stroma-free differentiation of hiPSCs into mature CD3⁺TCRαβ⁺ cytotoxic T lymphocytes (CTLs), overall efficiency remains limited. Here, we identify small-molecule modulators that enhance T cell output, particularly at the ProT cell stage. Targeted and stage-specific inhibition of AHR, DOT1L, or GSK3 drives robust maturation from ProT to CD4⁺ immature single-positive (ISP) cells, markedly increasing CD4⁺CD8⁺ populations and augmenting CTL production of up to 2000 fold. hiPSC-derived T (iT) cells matured under these conditions display superior activity in cytotoxicity assays using AMG-701 (BCMAxCD3) or Blinatumomab (CD19xCD3). These effects were reproducible across independent hiPSC lines, diverse hematopoietic progenitor generation methods, and multiple stroma-free differentiation platforms, and were further validated in cord blood CD34⁺ cells. Notably, AHR inhibition enhanced T cell development and promoted B lymphopoiesis, revealing shared regulatory pathways in lymphoid lineage specification. We also demonstrate that the Oct4-activating compound OAC1 functions as a weak AHR inhibitor, partially recapitulating the effects of canonical AHR blockers in both cellular and zebrafish AHR reporter systems. Collectively, our findings define key molecular circuits governing human lymphoid differentiation and establish practical strategies to optimize the yield and function of hiPSC-derived cytotoxic T cells. This work advances the development of both universal and autologous hiPSC-derived T cell therapies, offering a path forward even for patient-specific hiPSC lines with suboptimal T cell differentiation potential.

## INTRODUCTION

The recent successes with chimeric antigen receptor (CAR)-T cell immunotherapies offer hope for indications beyond hematologic malignancies, including solid tumors and auto-immune diseases (Choi et al., 2024; Mougiakakos et al., 2021). However, meeting this promise through conventional CAR-T cell therapy approaches alone faces significant challenges, such as the difficulty to scale-up manufacture of patient-autologous CAR-T cells, the possibility of production failure, and variable product activity (Abou-El-Enein et al., 2021) as well as the risk of causing product-derived T-cell malignancy that prompted the FDA to issue a black-label warning (https://www.fda.gov/vaccines-blood-biologics/safety-availability-biologics/fda-investigating-serious-risk-t-cell-malignancy-following-bcma-directed-or-cd19-directed-autologous). These considerations have galvanized the field of human induced pluripotent stem cell (hiPSC)- derived T cells. Given their scalability and amenability to precise genetic engineering of multiple loci, hiPSC-derived offer a compelling source for allogeneic T cells for use in T cell immunotherapy. Indeed, Fate Therapeutic’s FT819, a hiPSC-derived CAR-T cell product, has shown promising data in a Phase-1 clinical trial in patients with relapsed/refractory B cell lymphoma (https://clinicaltrials.gov/study/NCT04629729) and is now also being evaluated as a treatment for the auto-immune disease systemic lupus erythematosus (Kirkeby et al., 2025). Another FATE product, FT825 is currently in a clinical trial for solid tumors (https://www.cancer.gov/research/participate/clinical-trials-search/v?id=NCI-2024-01263).

Establishment of the hematopoietic system, commitment to lymphoid fate, and development into mature cytotoxic T cells are driven by sequential developmental phases, migration through distinct anatomical microenvironments, and complex regulatory mechanisms. T cell differentiation from hiPSCs is enabled by our still emerging understanding of the underlying stage-specific developmental cues and mechanisms. Following mesoderm and hemogenic endothelium specification, hiPSCs yield CD34^+^ hematopoietic progenitor cells (HPCs), with protocol-dependent outputs that differ in subsequent lymphoid potential. HPCs then further differentiate into ProT cells in the presence of Notch ligands and cytokines such as SCF, IL-7 and FLT3L (Chang et al., 2014; Jing et al., 2022; Mout et al., 2025). Early stage thymocytes are termed double-negative (DN) based on absence of both CD4 and CD8 expression. DN thymocytes develop into immature CD4 single–positive cells (ISP) that subsequently acquire CD8 expression, thereby becoming CD4/CD8 double–positive (DP) cells. *In vivo,* DP cells then develop either into mature, TCR**_αβ_**–expressing CD4 single positive (SP) helper T cells or into TCR**_αβ_** –expressing CD8-SP cytotoxic T cells, depending on interactions with either MHC class II or I molecules, respectively, on thymic epithelial cells. To date, most i*n vitro* differentiation protocols have a strong bias towards CD8SP maturation, with efficient CD4-SP T-helper cell induction being an area of ongoing investigation (Amirault et al., 2025; Jones et al., 2024; Kawai and Kaneko, 2026).

Although immune cells generated from iPSCs mirror those found in primary tissues, they typically retain distinct transcriptional profiles and functional attributes, and frequently present a phenotype that is either more primitive or skewed toward innate immune characteristics (Diop and van der Stegen, 2024). Further, differentiation protocols are lengthy and hampered by a great degree of cell death, in line with thymocyte selection *in vivo*, where the vast majority of DP progenitors are depleted through the combined mechanisms of neglect (lack of positive selection signals) and negative selection (removal of auto-reactive clones) (Klein et al., 2014; Scollay et al., 1980; Steier et al., 2024). Another shortcoming is the emergence of off-target cell types, like NK, NKT, and TCRγδ T cells (Jing et al., 2024; Li et al., 2023), the reasons for which remain poorly understood. Collectively, these challenges are impeding clinical translation of hiPSC-derived T cells. While therapeutic hiPSC CAR-NK cells are currently easier to produce and consequently are more advanced towards FDA approval (Diop and van der Stegen, 2024), their limited capacity for *in vivo* persistence and expansion will likely require co-treatment with CAR-T cells (Shimasaki et al., 2020), at least in the context of cancer therapy. Taken together, these shortcomings highlight the urgent need for further optimization of T cell differentiation protocols to fully harness their potential in clinical applications.

In this study, we identified several compounds through a live cell confocal-imaging based screening approach that substantially improves cell viability and efficiency of iT cell production while reducing the emergence of off-target, innate immune cells.

## RESULTS

### Small molecule screen identifies enhancers of T cell fate in hiPSC-derived ProT cells

We have previously demonstrated that culturing hiPSC-derived CD34^+^ HSPCs on DLL4 and VCAM1 in the presence of SCF, IL-7, TPO and FLT3L promotes acquisition of ProT cell fate from hiPSCs (Jing et al., 2022). However, subsequent maturation into ISP and DP cells, as well as specification of CD8-SP CTLs, represent major bottlenecks within this protocol and those published by others (Iriguchi et al., 2021; Jones et al., 2024). These bottlenecks can be partially overcome with lentiviral knock-down of EZH1 (Jing et al., 2022) a repressor of lymphoid fate (Vo et al., 2018). However, the variable efficacy of lentiviral knockdown combined with the risk of unpredictable adverse effects caused by insertional mutagenesis pose obstacles to translating this approach into safe and effective therapies. Chemical inhibition of EZH1 could be a viable alternative, but to date no EZH1-specific inhibitors have been described. In a recent study, we screened a library of epigenetic modulators and found that small molecule inhibition of G9A can compensate for the lack of lentiviral EZH1 knockdown. This demonstrates that pathways other than EZH1 can be leveraged to boost T cell yields (Jing et al., 2024).

Here, we assembled a bespoke library of small molecules (Supplemental Table S1), hand-picked to interrogate a broader range of targets beyond epigenetic factors, including many pathways relevant to hematopoietic and lymphoid cell development and regulation. We screened this library for agents capable of enhancing the cell survival, expansion, maturation of iT-proT cells into ISP cells while also limiting commitment to off-target lineages such as CD56^+^ NK cells (FIG 1A). Compounds were added to iT-day 14 cells, differentiated from hiPSC-derived CD34^+^CD43^-^ hemogenic endothelial cells (FIG S1A). At this stage, the cells are firmly committed to the lymphoid lineage, as evidenced by their uniform expression of the marker CD7^+^ and the emergence of CD7^+^CD5^+^ cells (FIG S1B), matching the profiles of proT1 and proT2 progenitor cell stages (Awong et al., 2009). Of note, we routinely detect presence of CD56^+^ cells at this stage as well (FIG S1B), consistent with the dual T/NK potential of proT1 and proT2 cells (Sánchez et al., 1994).

**FIG 1.**
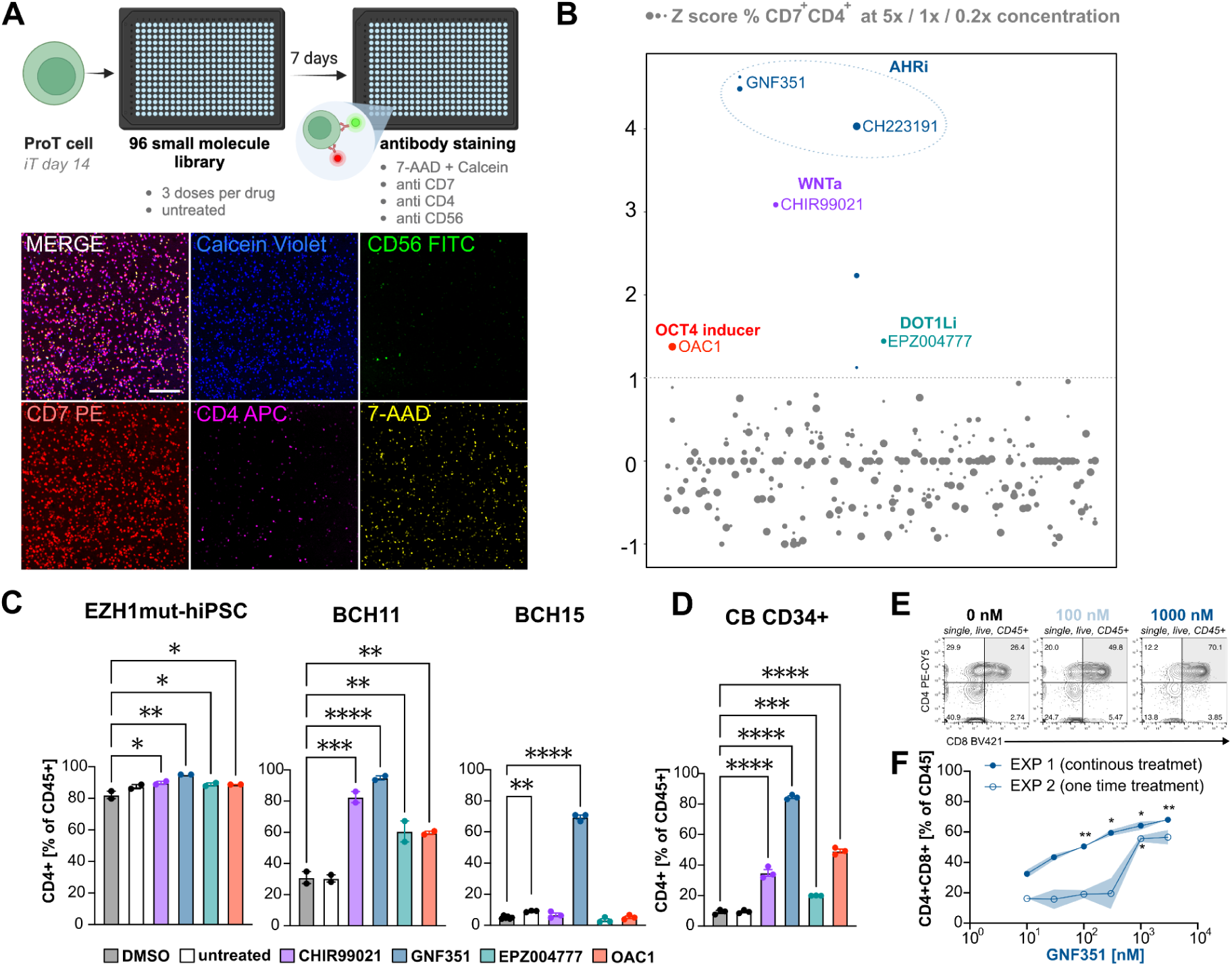
Small molecule screening identifies inducers T cell differentiation boosters. A: Schematic overview of the screening approach. hiPSC-derived CD34^+^ hemogenic endothelial cells were differentiated towards ProT cells for 14 days in a stroma-free protocol (Jing et al., 2022). At iT day 14 cells were harvested and replated into a 384 well plate and exposed to a small molecule library. After 7 days cells were stained with anti CD56, anti CD4, anti CD7 with 7-AAD and Calcein Violet viability dyes. Scale bar indicates 20 µm. B: Primary screening results. 5 hits were identified when a Z-score threshold of 1 was applied. C: Hit validation, bar graphs depicting mean percentage of CD4^+^ cells out of CD45^+^ population on iT day 28 in hiPSC line EZH1^mut/wt^ (screening line), BCH1157 and BCH1566. Error bars depict SEM of replicate wells, one way ANOVA comparison against DMSO. E: Exemplary flow cytometry plots on iT day 28 demonstrating an increased proportion of CD4^+^CD8^+^ DP cells after GNF351 treatment in BCH1157 hiPSC-derived ProT cells. F: Mean values plotted from two independent experiments with either continuous drug treatment (solid circles) or one time drug exposure (open circles). SEM of technical replicates is indicated by blue shading. Two-way ANOVA with Dunnett’s multiple comparison test against respective untreated cells.

Each compound was assayed at three different concentrations. Z-scores of compound-induced changes in the frequency of CD7^+^CD4^+^ cells were calculated for wells with cell viability above 50% based on Calcein violet positivity and exclusion of 7-AAD. We identified the Aryl hydrocarbon receptor pathway as the strongest candidate target, with the two highest-scoring agents (GNF351351 and CH223191) both being canonical AHR inhibitors (AHRi) (Kim et al., 2006; Smith et al., 2011) (FIG 1B). Besides increasing the frequency of CD7^+^CD4^+^ cells, both AHRi also induced a noticeable increase in total live cells. In contrast, presence of the gamma secretase inhibitor DAPT was not compatible with cell survival, emphasizing the expected continued Notch-dependence of T cell development at this stage of differentiation, and confirming the physiological behavior of the cells in our assay (FIG S1C). In addition, we followed up on all other hits: the WNT agonist CHIR99021 (3 µM), the DOT1L inhibitor EPZ004777 (1 µM), and OAC1 (5 µM) (‘OCT4 activating compound), an orphan compound originally identified as an inducer of OCT4 that was later shown to boost expansion of CB CD34^+^ HSPCs (Li et al., 2012).

Since our multi-color screening assay also assessed CD56 expression, we were able to likewise calculate the Z scores for each compound’s ability to induce the NK fate, which could be of value for developing effective manufacturing protocols for hiPSC-derived NK cell immunotherapies. Out of all compounds tested, the LSD1/CoREST-degrader UM171 (Subramaniam et al., 2020) had the strongest CD56-inducing effect (FIG S1C,D). UM171 has been previously described to increase the yield of hPSC-derived NK cells by up to 10-fold (Mesquitta et al., 2019). Its re-discovery in our screen therefore provides additional confirmation that our approach accurately models lymphopoiesis. We identified a total of 6 candidate NK fate boosting hits (FIG S1D) targeting multiple pathways, which will be explored in a separate study.

Many hiPSC differentiation protocols are optimized for specific, high-performing cell lines unique to individual labs. Consequently, these protocols often yield inconsistent results with other lines (Kattman et al., 2011; Maas et al., 2026), hindering reproducibility and complicating the transition to clinical applications, which require methods that work reliably with cGMP-grade line(s). To validate the T cell hits, we therefore derived ProT cells from three karyotypically normal hiPSC lines selected to cover a wide range of baseline T cell differentiation potential in the absence of hit compound addition. As a high baseline efficiency line, we used the EZH1^mut/wt^-hiPSC clone in which the original screen was performed. We also used its parental WT clone, line BCH1157 (Skorik et al., 2020). Lastly, we included the genetically unrelated hiPSC line BCH1566, a clone that despite robust differentiation in a lung epithelial cell differentiation protocol (Hawkins et al., 2021), fails to differentiate efficiently past the ProT cell stage, as evidenced by a lack of upregulation of CD4 (FIG S1E).

Interestingly, while all candidate hit compounds enhanced CD4^+^ ISP/DP cell production in EZH1^mut/wt^ and BCH1157 hiPSCs, only AHR inhibition (GNF351, 1 µM) induced a statistically significant improvement in iT cell differentiation in BCH1566 (FIG 1C). To extend these findings beyond hiPSC-derived progenitors, we next evaluated the effects of the compounds on T cell differentiation started from CB CD34^+^ HSCs that have a known high intrinsic potential to produce lymphoid cells (De Smedt et al., 2011). Similar to the results obtained with the EZH1 mutant line and BCH1157, addition of any one compound led to significant increases in the frequencies of CB derived CD4^+^ cells (FIG 1D). Of note, with all cell sources, addition of the AHR inhibitor GNF351 had the strongest effect.

In most cases, a single dose addition of GNF351 (1 µM) on iT day 14 was sufficient to increase CD4^+^ cell frequencies and yields, while BCH1566 cells displayed a stronger effect when the compound was added during each half media change (FIG S1F). We next wanted to test if the positive effect of AHR inhibition could be reproduced in the context of an alternative, stroma-free T cell differentiation protocol based on a commercial media or when the starting population of CD34^+^ cells was derived by a recently published protocol (Fowler et al., 2024) that more efficiently generates HPCs expressing medial HOXA genes and HLF, genes commonly associated with definitive myelo-lymphoid potential (Calvanese et al., 2022). The StemSpan T cell generation kit from Stem Cell Technologies follows a similar approach to the protocol developed by our group (Jing et al., 2022) using SFEMII medium supplemented with either lymphoid expansion or maturation supplement in a stage-dependent manner (FIG S1G). GNF351 treatment again significantly improved the yield and frequency of CD4^+^CD8^+^ cells irrespective of which hematopoietic progenitor production method or T cell differentiation process was used (FIG S1H). When we tested different GNF351 concentrations, we observed a highly dose-dependent increase in CD4^+^CD8^+^ double positive cells, with 100 nM being sufficient for a significantly improved DP cell production when cells were continuously exposed to the AHR inhibitor. With one-time drug addition, a higher dose of 1000 nM was necessary to promote formation of DP cells (FIG 1E,F). Taken together, we have identified AHR activity as a strong repressor of *in vitro* T cell maturation, with chemical inhibition of AHR facilitating acquisition of CD4^+^CD8^+^ double positive fate from ProT cells independent of the source, production method, or genetic background of the input cells or the T cell derivation method used, underscoring the robustness and reproducibility of our findings.

### AHRi promotes T and B cell differentiation and suppresses NK cell differentiation

To further investigate the role of AHR during ProT cell differentiation, we exposed CB CD34^+^ derived ProT cells to two structurally diverse AHR inhibitors (GNF351, CH223191) and two AHR agonists (FICZ (Rannug et al., 1995), BNF (Poland and Glover, 1980)). Agonists and inhibitors had consistently opposite effects on iT cell differentiation outcomes, with AHR inhibitors drastically increasing the number of CD45^+^ cells, as well as the percentage of DP-iT cells and the frequency of CD3^+^-iT cells, while drastically decreasing formation of off-target CD56^+^ innate immune cells (FIG 2A). When those cultures were differentiated further, it became evident that both AHR inhibitors increased the number of CD3^+^CD56^-^ T cells, with CH223191 additionally increasing the number of CD3^+^CD56^+^ NKT cells (FIG 2B). We observed a similar skewing favoring adoptive T cell development over innate NK cell fate upon AHR pathway inhibition when treating hiPSC-derived ProT cells, indicating an inverse correlation between AHR pathway activity and T cell fate choice (FIG 2C, FIG S2A).

**FIG 2.**
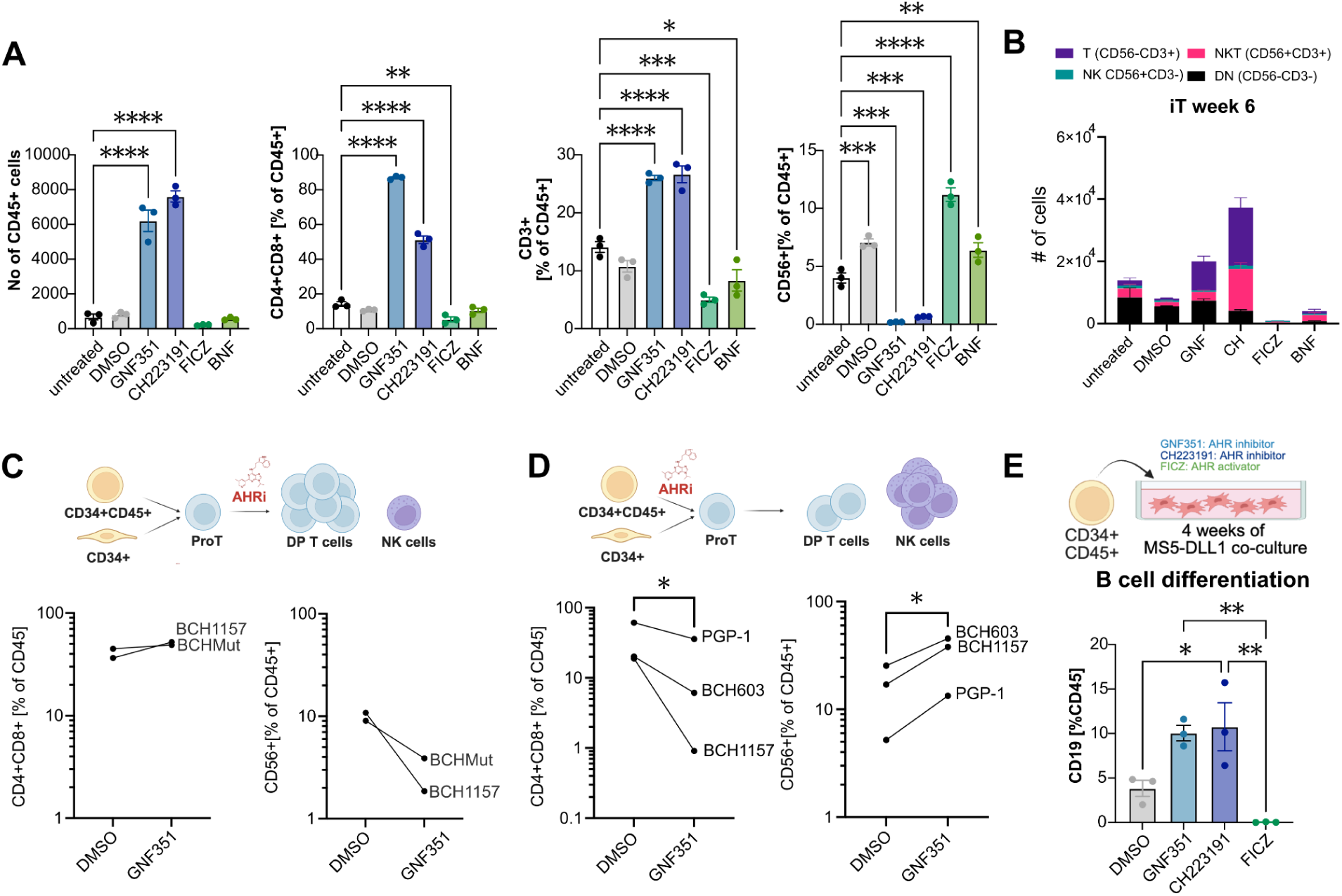
AHR inhibition promotes pan lymphoid potential. A: AHR inhibitors GNF351 and CH223191 both increase number and percentage of CB CD34^+^-derived CD4^+^CD8^+^ cells at the expense of CD56^+^ cells, while AHR activators FICZ and BNF have the opposite effect. B: Distribution of T (CD56-CD3^+^), NKT (CD56^+^CD3^+^), NK (CD56^+^CD3^-^) and double negative (CD56^-^CD3^-^) cells in week 6 T cell differentiation cultures. C: Schematic depicting T cell vs NK cell differentiation outcome, depending on timing of AHR inhibition and flow cytometric assessment of T vs NK cell differentiation in two hiPSC lines after AHR inhibition (GNF). D: Flow cytometric assessment of T vs NK cell differentiation in three hiPSC lines after AHR inhibition (GNF) prior to the ProT cell stage. Paired t-test p=0.014 and 0.029, respectively. E: Schematic and flow cytometric assessment of B cell differentiation outcome after 4 weeks of CB CD34^+^ culture with AHR inhibitors GNF351 or CH223191 or AHR agonist FICZ. Bar graphs depict the mean and SEM of 3 independent CB donors. One way ANOVA with Tukey’s multiple comparison test.

Interestingly, the T cell–promoting effect is highly dependent on the treatment window: when hiPSC-derived CD34⁺ cells are exposed to GNF351 before the ProT cell stage and then carried through our remainder of our T cell production protocol, GNF351 reduces DP cell yields and instead preferentially augments CD56⁺ cell production (FIG 2D). These data reveal a biphasic, time-dependent role for the AHR pathway during T lymphocyte development, where modulating agents exert opposing effects at distinct developmental stages.

Next, we asked whether AHR inhibition promotes T cell fate specifically, or whether the effect extends to other lymphoid lineages. To this end, we assessed the effect of AHR pathway modulation during B cell directed *in vitro* differentiation of CB derived CD34^+^ HSCs. In the presence of either AHR inhibitor (GNF351351 or CH223191), cells displayed increased differentiation towards CD19^+^ lymphocytes, while the AHR agonist FICZ completely ablated B cell differentiation potential (FIG 2F, S2B). These data indicate that AHR inhibition improves adaptive immune cell differentiation at the expense of innate immune cells.

### Dynamic requirement for WNT activation during T cell differentiation

WNT is known to play a crucial role during *in vivo* T cell maturation, regulating thymocyte development and proliferation (Schilham et al., 1998; Staal et al., 2001; van de Wetering et al., 2002). This prompted us to test if earlier administration of CHIR99021 would further improve iT differentiation in our protocol. Although WNT activation at the ProT cell stage has a positive effect, this benefit is highly stage-dependent, as administering CHIR99021 during the first two weeks of iT cell culture negatively affected ProT cell differentiation (FIG S1C). The fact that we observed a stronger positive effect on T cell differentiation when CHIR was continuously administered from day 14 onwards as opposed to single drug exposure on day 14, led us to speculate that WNT activation is mostly beneficial at later stages of iT differentiation in our system. Indeed, we found that including CHIR99021 at the CD4^+^CD8^+^ double positive stage drastically improved differentiation to the mature CD3^+^TCRɑβ^+^CD4^-^CD8^+^ (‘CD8 single positive’) stage of T cell development. This observation was validated across two hiPSC lines (FIG S2D).

### Transcriptional profiling of small molecule treated ProT cells

To assess immediate transcriptional downstream effects of small molecule exposure, we exposed CB CD34^+^ derived day 14 ProT cells from two independent donors for 24h to our screening hits, and performed bulk RNA sequencing of two replicates each. Batch correction was performed to account for the significant donor-dependent variability: compare sample clustering pre- (FIG S3A) and post-correction (FIG S3B). RNA-seq data confirmed expression of canonical Pro T cell markers such as CD7, CD5, CD44, and *BCL11B* as well as absence of CD56/*N-CAM1* (FIG S3C). Of note, CD44 expression levels were higher in CHIR-exposed cells, in line with proposed connections between WNT signaling and CD44 expression (Walter et al., 2022; Wielenga et al., 1999). Short term exposure of cells to EPZ004777 barely induce transcriptional changes (19 downregulated, 11 upregulated genes, FIG 3A), which is expected given that it can take several days before EPZ004777 treated cells show a significant loss in H3K79 methylation (Kurani et al., 2022). Conversely, the canonical AHR target gene AHRR (Baba et al., 2001) was strongly downregulated in GNF351 treated cells (FIG 3B). Another strongly down-regulated gene, CYP1B1, is also a direct target of AHR (Yang et al., 2008), confirming successful inhibition of the AHR pathway in GNF351 treated samples (FIG 3B). While OAC1 treatment induced very few transcriptional changes, AHRR and CYP1B1 both were among the strongest down-regulated genes, indicating that this orphan compound might antagonize the AHR pathway similarly to GNF351 (FIG 3B). While their precise downstream mechanisms remain to be elucidated, CHIR, GNF351 and OAC1 all upregulated expression of LYL1, a transcription factor with a known role in lymphoid specification and maintenance of early T cell progenitors (Zohren et al., 2012), as well as WNT target gene TCF7 (Roose et al., 1999), which encodes the transcription factor T cell factor 1 (TCF-1), further validating these compounds as T cell differentiation promoting agents (FIG S3C).

**FIG 3:**
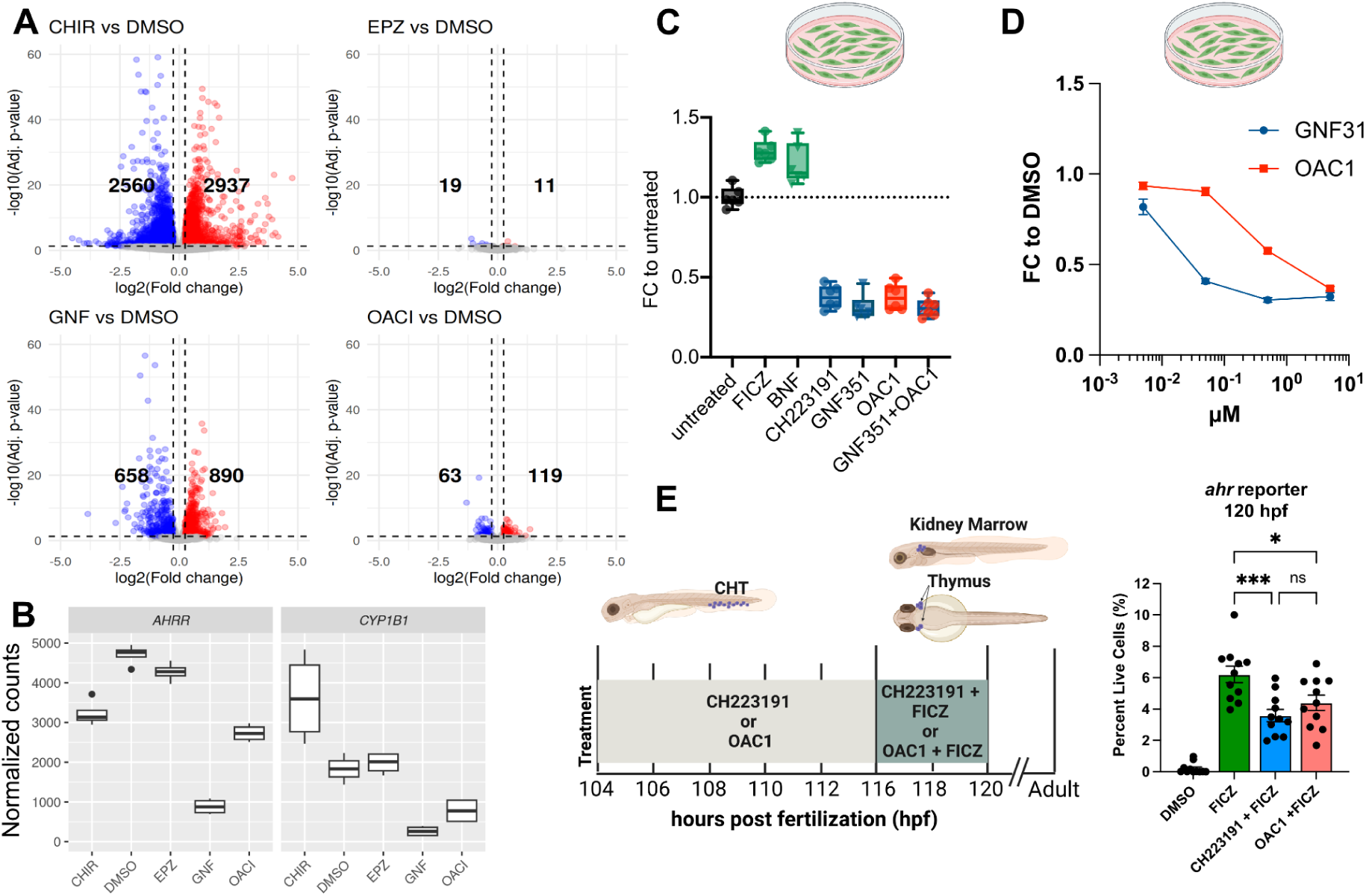
OAC1 is a weak AHR inhibitor. A: RNA-seq analysis of CB CD34^+^ differentiated towards ProT cells and exposed to indicated drugs for 24h. Volcano plot indicating number of differentially expressed genes (DEGs) between drug treated vs DMSO ctrl samples. B:Expression level of AHRR and CYP1B1 are most strongly repressed in cultures treated with the known AHR inhibitor GNF351 but also in cultures treated with OAC1 C: FICZ and BNF increase expression of GFP in AHR reporter 293T cells, while CH223191, GNF351 and OAC1 decrease GFP expression relative to DMSO. D: Mean fluorescent intensity (MFI) of GFP in AHR reporter cells treated with increasing doses of GNF351 or OAC1. Mean +/- SEM. E: Both CH223191 (1 µM) and OAC1 (5 µM) are able to inhibit FICZ (0.2 µM) driven activation of the *ahr* activity reporter in zebrafish. Each dot represents a fish, the experiment has been performed three independent times.

### OAC1 phenocopies AHR inhibition

Next, we sought to confirm that exposing cells to OAC1 indeed inhibits the AHR pathway, either directly or indirectly. To this end, we stably introduced an AHR reporter into 293T cells, in which GFP-expression is regulated by the promoter of the CYP1A1 gene, another canonical AHR target gene (Weems and Yost, 2010). We tested the reporter line by exposing it to known AHR activators (FICZ, BNF) as well as inhibitors (CH223191, GNF351), which resulted in the expected changes in GFP expression, thus validating this system as a faithful reporter of AHR activity. Exposing reporter cells to OAC1 also decreased the percentage of GFP^+^ cells relative to DMSO ctrl treated cells, supporting our hypothesis that OAC1 exposure results in AHR pathway inhibition (FIG 3C). Of note, there was no additive effect of OAC1 in combination with GNF351 (p=0.9121), suggesting that OAC1 does not function through a different mechanism to reduce GFP expression (FIG 3C).

Next, we treated reporter cells with increasing concentrations of either GNF351 or OAC1 followed by flow cytometric quantification of GFP. OAC1 treated cells displayed a dose-dependent reduction in GFP fluorescence, much like what we observed with GNF351, albeit requiring a higher dose indicating that OAC1 is a less potent AHR antagonist or perhaps acts as an indirect inhibitor of this pathway (FIG 3D). This hypothesis was further tested *in vivo* using a transgenic ahr activity reporter zebrafish line. In this line, GFP expression is placed under the control of a promoter with eight AHR binding sites, allowing us to read out AHR activation via fluorescent signal detection. Reporter-transgenic embryos were first exposed to CH223191, a known AHR inhibitor and validated hit in our screen, serving as a positive control in this assay, or to OAC1 at 106 hours post-fertilization. Then, these embryos were dosed with the AHR agonist FICZ and the antagonists refreshed at 116hpf followed by analysis at 120hpf. Canonical AHR inhibitors such as CH223191 will bind the receptor and block transcription of GFP even in the presence of activator FICZ (Mohammadi-Bardbori et al., 2019). The number of GFP positive cells was significantly decreased in the presence of CH223191 compared to the activator alone. A similar – albeit less dramatic – decrease was observed in embryos exposed to OAC1, further supporting the notion that OAC1 has direct or indirect AHR inhibitory function (FIG 3E).

### DOT1L inhibitor EPZ004777 acts synergistically with AHRi to boost iT cell differentiation

Next, we interrogated whether the identified hits would work synergistically. Given that OAC1 appears to largely phenocopy AHR inhibitors, we speculated it would not further improve iT cell differentiation when combined with GNF351, the most potent AHRi and T cell development-promoting agent identified in our study. In contrast AHRi, WNTa and DOT1Li likely act through distinct pathways, which could potentially synergize to maximize iT cell differentiation outcome. Pro T cells from two independent hiPSC lines (BCH1157, BCH1566) were treated either with GNF351 alone or GNF351 in combination of either WNTa or DOT1Li or both. Indeed, combination of AHRi (GNF351) with DOT1L inhibition (EPZ004777) further enhanced acquisition of a CD4^+^CD8^+^ phenotype in week 4 cultured cells: the effect was highly significant for hiPSC line BCH1566 and showed a positive trend for BCH1157 (p=0.073). Acquisition of CD4^+^CD8^+^ was not further improved by inclusion of WNTa (FIG 4A).

**FIG 4:**
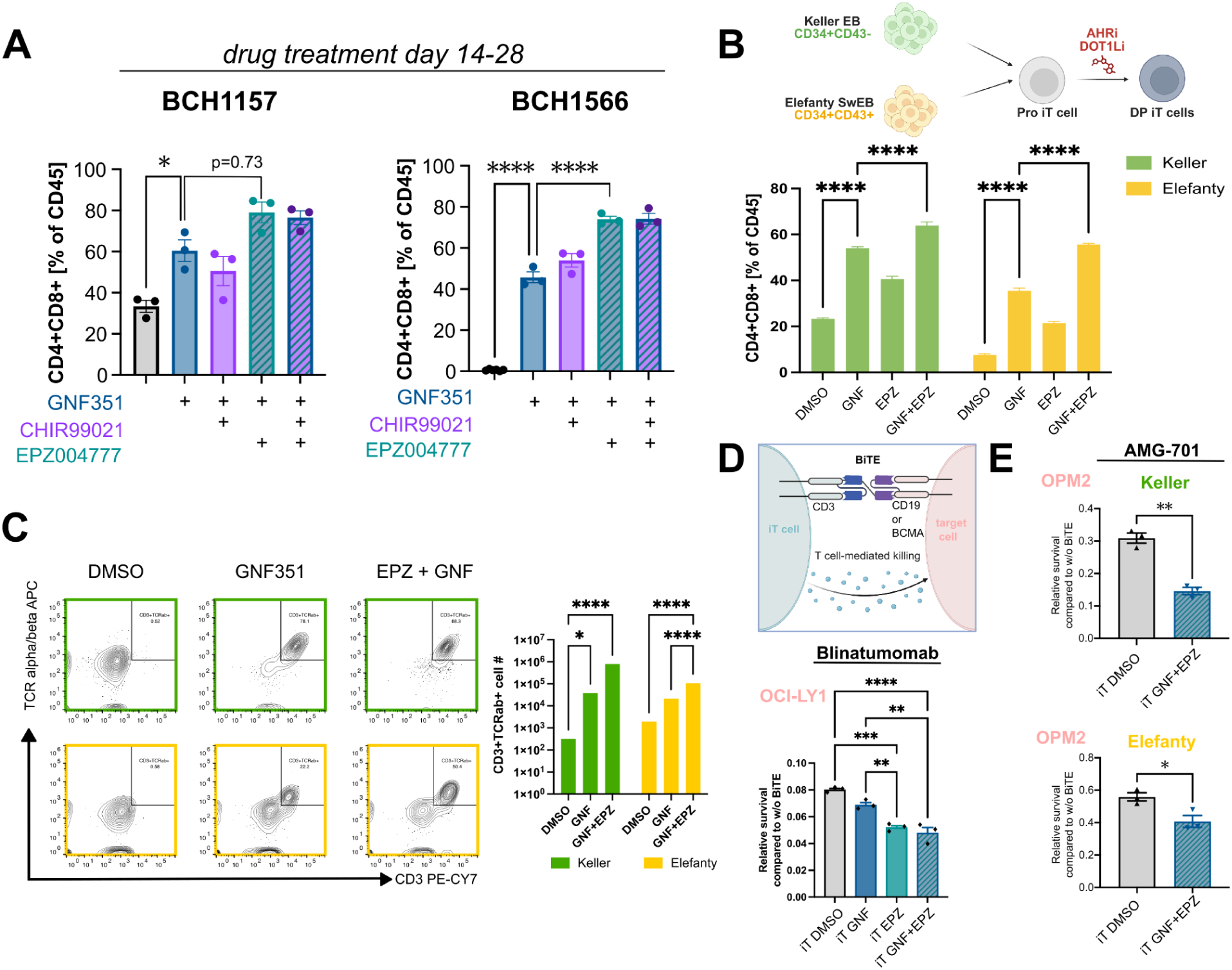
AHRi and DOT1Li synergistically improve iT cell differentiation outcome and cytotoxicity. Flow cytometric profiling of week 4 iT cells derived from two hiPSC lines treated with indicated drug combinations from iT day 14 onwards. A GNF351 and EPZ004777 synergistically increase the percentage of CD4^+^CD8^+^ DP cells in two independent hiPSC lines. B: Percentage of CD4^+^CD8^+^ cells after iT differentiation of either Keller- or Elefanty-derived hematopoietic cells in the presence of small molecule inhibitors. C: Representative flow plots and quantification of CD3^+^TCRɑ□^+^ cells post single positive induction with Immunocult anti CD3/CD28 antibodies. D: Relative survival of target cells (OCI-LY1) to no BiTE ctrls after 48h of co-culture with iT cells derived in the presence of small molecule inhibitors. Blinatumomab was added at 1 ng/ml, E:T of 5:1, One-way ANOVA E: Relative survival of target cells (OPM2) to no BiTE ctrls after 48h of co-culture with iT cells derived in the presence of small molecule inhibitors. Keller and Elefanty indicate which HPC protocol was used. Mature iT cells were pre-sorted for live, CD8^+^CD56^-^ prior to setting up the co-culture. AMG-701 was added at 3nM, E:T of 1:1, unpaired t-test. * indicates p≤ 0.05. **P≤0.01

### A Distinct Embryoid Body Differentiation Protocol Further Enhances iT cell Yields

While the data presented above was generated with hematopoietic progenitors differentiated following an embryoid-body (EB) based protocol defined by Keller and colleagues (Ditadi et al., 2015), improved protocols for more efficient production of HPCs with lymphoid potential have recently been published by the Loh and Elefanty laboratories (Fowler et al., 2024; Ng et al., 2024; Zheng et al., 2025). The Keller protocol entails deriving iT cells from CD34^+^CD43^-^hemogenic endothelium isolated from day 8 dissociated EBs. In contrast, the Elefanty protocol generates highly uniform populations of CD43^+^CD34^+^ HPCs and HSC-like cells that have already undergone endothelial-to-hematopoietic transition, thus removing the need for MACS-based enrichment (FIG S4A). Of note, the Elefanty protocol generates ProT cells at a much higher frequency, resulting in a 1000x greater yield per input hiPSC compared to the Keller protocol (FIG S4C). This is partially due to greater expansion between iT day 0 to 14 compared to cells produced using the Keller protocol, with Elefanty protocol derived cells having nearly the same potential as primary CB CD34^+^ cells in the same protocol (FIG S4C).

To determine whether the hit compounds identified in our chemical screen exert consistent effects regardless of the hematopoietic differentiation method, we performed side-by-side iT cell differentiation using HPCs generated from the same hiPSC line via both protocols (Keller vs Elefanty (Ng et al., 2024)). Consistent with our prior results, GNF351 treatment alone and in combination with EPZ004777 significantly increased the percentage of DP cells in both cases (FIG 4B), and could be robustly replicated (FIG S4D). In both instances, the double drug treatment resulted in significantly higher DP cell frequencies compared to the no- or single-compound conditions (FIG 4B). Importantly, small molecule treatment after the ProT cell stage of differentiation also resulted in dramatically increased CD3^+^TCRɑβ^+^cell yields (FIG 4C), which represents a fold increase of 2253 +/- 10 over DMSO controls in case of iT cells following the Keller protocol and 57 +/- 13 fold increase for Elefanty, respectively.

### Chemically-derived iT Cells Mediate Enhanced Cytotoxicity

Next, we utilized bi-specific T cell engagers (BiTE) to assess the cytotoxic potential of *in vitro* differentiated iT cells derived in the presence of either GNF351 alone, EPZ004777 alone or a combination of both. Differentiated iT cells were co-cultured with BiTE-directed target cells (OCI-LY1 for Blinatumomab CD19xCD; OPM2 for AMG-701 BCMAxCD3). In these *in vitro* killing assays iT cells differentiated in the presence of small molecules mediated significantly greater cytotoxicity than DMSO control iT cells, with cells derived using both inhibitors consistently exhibiting the most robust killing activity (FIG 4D), even when using Odronextamab (CD20xCD3) to target Raji cells, which were more resistant to killing (FIG S4F).

To confirm specificity of lysis, we used K-562 with and without CD19 overexpression as target cells for Blinatumomab mediated iT cell killing. Importantly, iT cells differentiated with GNF351 and EPZ004777 showed no baseline cytotoxicity against CD19 negative cells (K-562), comparable to CD8 PB T cells. DMSO ctrl iT cells showed comparable cytotoxicity towards K-562 regardless of CD19 overexpression, likely due to NK-mediated cytotoxicity in line with the higher percentage of CD56 positive cells in cultures differentiated without small molecule addition (FIG S4G).

## Discussion

In this study, we identified AHR inhibitors (GNF351, CH223191) as potent and stage-specific effectors that significantly enhance the maturation of hiPSC-derived Pro T cells into CD4^+^CD8^+^ double positive T cells. This discovery addresses a critical challenge in immunotherapy: the efficient and scalable generation of functional T cells from pluripotent stem cell sources. By leveraging confocal immunofluorescence imaging, we uncovered 5 hit compounds that consistently promoted T cell lineage commitment and maturation, as confirmed by flow cytometric based immunophenotyping. These findings provide novel tools for improving *in vitro* T cell production and offer valuable insights into the molecular pathways governing hematopoietic and lymphoid differentiation. Our screening results indicate that the AHR pathway serves as a critical regulator at the intersection of T cell and NK cell lineage commitment, with an active pathway promoting NK cell fate, while inhibition favors T cell fate (and under appropriate culture conditions improves B lymphocyte development).

The two key advances of our small molecule-enhanced iT cell differentiation protocol are the entirely transgene-free design (compared to the EZH1 knockdown-dependent protocol;(Jing et al., 2022) and the robustness across genetic background, input cell sources, and T cell differentiation methods. This protocol consistently generates mature CD3⁺TCRαβ⁺ T cells across diverse hiPSC lines, including those that previously failed during iT cell maturation, thus making it well-suited for translational efforts.

The two key advancements of our small molecule-enhanced iT cell differentiation protocol are its entirely transgene-free design compared to the EZH1 knockdown dependent protocol (Jing et al., 2022) and its significantly improved robustness across genetic background, input cell sources, and T cell differentiation methods. This protocol consistently generates mature CD3⁺TCRαβ⁺ T cells across diverse hiPSC lines, including those that previously failed during iT cell maturation stages, thus making it well-suited for translational efforts.

As we curated the small molecule library with carefully selected compounds likely involved in definitive hematopoiesis, developmental regulators, and lymphopoiesis, most of the hits are modulators of well-established pathways, such as AHR and WNT. Originally identified in an unbiased screen, AHR inhibitors are well known to promote *ex vivo* expansion of HSPCs (Boitano et al., 2010). The observation of increased numbers of bone marrow-resident Lin-Sca^+^Kit^+^ HSPCs in *Ahr* knockout mice revealed the importance of the AHR pathway during vertebrate hematopoietic development *in vivo*(Singh et al., 2011). Additional important roles for AHR in regulating erythropoiesis and megakaryopoiesis have since been described (Smith et al., 2013). Furthermore, overexpression of a constitutively active AhR in mouse thymocytes led to a decreased number of thymocytes and an increase in CD8 single-positive cells (Nohara et al., 2005). StemRegenin 1 (SR-1), an AHR inhibitor, has demonstrated potential in *ex vivo* expansion of HSCs. However, it has not yet received FDA approval. The regulatory status of SR-1 remains uncertain, as a clinical trial evaluating the efficacy of expanded umbilical cord blood CD34^+^ cells using this compound was discontinued (clinicialtrials.gov, ID NCT02765997). A previous study assessing the effect of AHR inhibition on early hematopoietic differentiation found that it improves the development of conventional NK cells from hESCs whereas AHR hyperactivation favors development of group 3 innate lymphoid cells (ILC3s) (https://www.ncbi.nlm.nih.gov/pmc/articles/PMC5492090/.

While the effects of AHR modulation on immature HSPCs, non-lymphoid hematopoiesis, and mature T cell regulation have been well characterized, the role of AHR signaling during early lymphoid progenitor differentiation was previously unknown. In our screen, AHR inhibition had the most robust phenotype of all small molecule hits and was able to significantly increase CD4^+^CD8^+^ output. Underscoring the reproducibility and generalizability of this connection, we were able to observe similar effects independent of the genetic background of the hiPSC line, their baseline T cell differentiation potential, the method through which the hiPSC-derived CD34^+^ cells were generated, the origin of the CD34^+^ HPCs (cord blood or hiPSC-derived), or which T cell differentiation medium was used.

Our discovery that the orphan hit compound OAC1 (OCT4 activating compound 1) may act as an inhibitor of the AHR pathway for the first time proposes a specific molecular mechanism to explain its activity. We derived our hypothesis that OAC1 primarily acts as a novel AHR inhibitor through multiple independent lines of evidence, including up-regulation of canonically AHR-repressed target genes in OAC1-exposed cells, repression of AHR activity reporter genes in cultured human cells and transgenic zebrafish embryos, and abrogation of OAC1 mediated effects on cells in the presence of saturating amounts of GNF351. Of note, our hypothesis is consistent with all previously described activities of OAC1. For example, the effects of OAC1 on human HPCs are very similar to those observed with canonical AHR inhibitors, with both approaches promoting HPC expansion and HOXB4 induction (Boitano et al., 2010; Huang et al., 2016; Jiang et al., 2018; Wagner et al., 2016; Zhu et al., 2021). In another context, OAC1 was originally discovered as an inducer of OCT4 expression during acquisition or maintenance of pluripotency (Li et al., 2012). Other studies focusing on the effects of AHR pathway modulation on pluripotency and OCT4 expression yielded similar results, linking AHR inhibition with increased OCT4 expression (Cheng et al., 2015; Nacarino-Palma et al., 2021). Of note, we saw no evidence of OCT4 induction by OAC1 in our T cell protocol, further supporting the notion that OAC1 acts as an AHR inhibitor with context-dependent outcomes as opposed to it functioning as a direct inducer of OCT4.

It is perhaps not surprising that the WNT agonist (WNTa) CHIR99021 emerged as a top hit in our screen, given the extensive prior evidence that WNT pathway inhibition - via soluble Frizzled decoys, DKK1, or cell-autonomous inhibition of β-catenin/TCF complexes - disrupts thymocyte development at multiple stages *in vivo*. More unexpected was the narrow temporal window for WNT activation we observed: premature WNTa exposure impaired progression to the ProT stage, whereas later activation enhanced DP and SP output. This is seemingly at odds with the requirement for TCF-1 from the earliest stages of T cell lineage specification throughout positive and negative selection in the thymus (Shin et al., 2024). A likely explanation for this apparent discrepancy is that our *in vitro* system collapses multiple spatially and temporally segregated thymic cues into a single, chemically defined environment. In the thymus, WNT ligands, Notch signals, and cytokines are presented in distinct niches and gradients, and TCF-1 integrates both canonical WNT and WNT-independent inputs in a stage-specific manner (Weber et al., 2011). By contrast, global activation of WNT/β-catenin with CHIR99021 on pluripotent- or HE/HP-stage cells may reinforce stemness programs and alter Notch responsiveness, thereby delaying or diverting commitment toward the ProT fate, even though later WNT activity remains beneficial for thymocyte survival and maturation. Thus, early CHIR99021 exposure likely perturbs the timing and context of WNT signaling rather than contradicting the fundamental requirement for TCF-1 during early T cell development (Ma et al., 2012).

Consistent with this, one mechanistic hypothesis is that WNTa at the maturation stage primarily acts on DP thymocytes by inducing the pro-survival molecule BCL-XL, as previously shown in mouse thymocytes (Taylor and Rudd, 2017). In addition, stabilized β-catenin has been reported to modulate positive selection and IL-7 receptor signaling, resulting in increased CD8 SP thymocyte output (Yu et al., 2007). Together, these findings support a model in which canonical WNT signaling is required throughout T cell development but is normally gated by niche-specific cues; in our *in vitro* system, CHIR99021 uncouples this pathway from its physiological spatiotemporal constraints, creating an early window in which WNT activation is detrimental to ProT specification yet a later window in which it is advantageous for DP and CD8 SP survival and maturation. To our knowledge, CHIR99021 has not previously been incorporated into the ProT-to-T cell phase of human PSC-derived T cell differentiation protocols, despite its widespread use at earlier stages to promote definitive hematopoiesis (Kennedy et al., 2012).

Our last screening hit EPZ004777 is an inhibitor of mammalian disruptor of telomeric silencing 1 like protein (DOT1L). DOT1L is a histone three lysine 79 methyl transferase implicated in early embryonic development and normal hematopoiesis, as DOT1L knockdown leads to impaired blood lineage formation (Arnold et al., 2022). Interestingly, prior evidence suggests that DOT1L inhibition could be beneficial for early T cell differentiation, as DOT1L knock-down was a primary hit promoting CD4^+^CD8^+^ cell fate in an shRNA screening against epigenetic modifiers. However, subsequent validation experiments did not demonstrate statistically significant efficacy, indicating that the effects observed in the primary screen of that study were not reproducible under more stringent testing conditions (Vo et al., 2018).

While shRNA mediated knockdown and small molecule inhibitions are different modes of action, our data would indicate that the primary hit was not a false positive but rather that DOT1L inhibition is context specific or has a modest effect size and its positive effect on T cell differentiation needs to be further investigated. It is particularly intriguing that DOT1L has a synergistic effect with AHR inhibition, indicating that we are far from having fully understood *in vitro* lymphocyte development and there are multiple pathways, whose modulation can further increase target cell output. Interestingly, DOT1L has been postulated to play a central role in CD8^+^ T cell differentiation by preventing premature antigen-independent differentiation and maintaining epigenetic integrity (Kwesi-Maliepaard et al., 2020). In light of these data and the fact that in our study DOT1L inhibition has been performed at the ProT cell stage, later exposure to DOT1L inhibitor EPZ004777 should be carefully evaluated.

The small molecules identified in this study are particularly impactful for hiPSC lines that exhibit minimal or absent baseline capacity to generate mature CD3⁺TCRαβ⁺ T cells under standard conditions, a challenge that mirrors the well-recognized variability in differentiation potential across hiPSC lines and donors (Kattman et al., 2011; Maas et al., 2026). We observed that specific patient-derived lines were intrinsically resistant to standard T cell differentiation. Strikingly, treatment with these compounds overcame this developmental block, not only enabling robust CD3⁺TCRαβ⁺ maturation but also actively suppressing alternative cell fates.. This pharmacologic intervention yields a defined, T cell-enriched product by simultaneously rescuing T cell output from non-productive lines and restricting off-target differentiation. Crucially, the robustness of our protocol across multiple hiPSC lines directly addresses a major hurdle in clinical translation. The pool of fully qualified, cGMP-grade hiPSC clones is often limited, and evaluating multiple lines to find an efficient differentiator is both time-consuming and cost-prohibitive. Because both autologous and allogeneic manufacturing workflows must maximize the utility of these valuable starting materials, differentiation protocols must be broadly effective. By lowering batch failure rates and ensuring successful lineage commitment regardless of baseline clone efficiency, this approach provides a highly feasible and regulatory-friendly framework for clinical manufacturing.

## METHODS

### hiPSC culture

For maintenance culture, hiPSCs were grown on Matrigel-coated dishes in Stemflex medium (ThermoFisher Scientific). Cells were routinely clump passaged using ReleSR (StemCell Technologies) when reaching ∼70% confluency. The EZH1^mut/wt^-hiPSC line was derived from BCH1157-hiPSCs via CRISPR/Cas9 targeting, resulting in heterozygous editing consisting of a 7 nucleotide insertion in the EZH1 locus, causing alternative, in-frame splicing and subsequent loss of 3 amino acids (ΔGEE).

### Hematopoietic progenitor differentiation

CD34^+^CD43^-^ hemogenic endothelial cells were differentiated from hiPSCs following a modified protocol developed in the Keller lab (Ditadi et al., 2015; Lummertz da Rocha et al., 2022). As an alternative, HSPCs (CD34^+^CD43^+^) were differentiated using the recently published protocol developed in the Loh lab (Fowler et al., 2024), as well as following the Swirl EB method developed in the Elefanty lab (Ng et al., 2024). hiPSC line BCH1157 has been previously published (Jing et al., 2022, 2024; Lummertz da Rocha et al., 2022; Skorik et al., 2020).

### T cell differentiation

T cell differentiation was performed with hemogenic endothelial cells differentiated from hiPSCs following the protocol developed by the Keller laboratory unless otherwise stated. CD34-Magnetic-activated cell sorting (MACS) purified day 8 EB derived-CD34^+^ hemogenic ECs (1e+5) were seeded on individual wells of a non-TC treated 24-well plate coated with DLL4-Fc (10 µg/ml) and VCAM-1 (2 µg/ml) for 2h at room temperature or at 4C overnight. T cell differentiation was performed following the protocol developed by Jing et al. without EZH1 lentiviral knockdown (Jing et al., 2022). In addition, to generalize the effect of the small molecules, CB derived CD34^+^ cells or various hiPSC-derived hematopoietic progenitors derived from 2D cultures (Fowler et al., 2024) or Swirl-EBs (Ng et al., 2024) were differentiated into iT cells following the same protocol. Of note, due to the different lymphoid differentiation potential of those various HSPC sources, plating cell densities had to be adjusted (5e+3 for CB CD34^+^ and Elefanty derived HPCs and 5e+4 for Loh-derived HPCs). In some cases, cells were differentiated in StemSpan™ SFEM II (Catalog # 09655) supplemented with StemSpan™ Lymphoid Progenitor Expansion Supplement (Catalog # 09915), followed by culture in StemSpan™ SFEMII supplemented with StemSpan™ Cell Progenitor Maturation Supplement (Stem Cell Technologies, Catalog # 09930).

### Small molecule screening for T cell inducers

Each compound was assayed at three different concentrations (at about 5x, 1x and 0.2x, with 1x equivalent to a commonly used dose for the compound). The library compounds were reconstituted and stored at -30°C in sealed in polypropylene microtiter plates (Abgene, #1055) containing 10µl per well (see also Suppl. Table S1). Rows A-O of a glass bottom 384-well plate (Cellvis, #P384-1.5H-N) were coated with DLL4 (Thermo Fisher) and VCAM1 by adding 25 µl/well with a 300 µl 12-channel pipette followed by spinning in a swing-out bucket rotor for 5 min at 300g and incubation in a cell culture incubator for 1h at 37 °C. The coating solution was removed by upside-down centrifugation into a receptacle (1 min. at 1,000g) followed by washing with 70µl PBS-/- and plating of 5,000 day 14 iT cells (EZH1^mut/wt^-hiPSC) in 50µl of medium per well. One µl of compound was added to a 96 well U-bottom plate containing 57 µl medium. After mixing, 8.1µl were added to quadrant 1 (1:416 dilution) of the 384 well plate followed by mixing and serial transfer of 10µl through quadrants 2 and 3 (1:2.5k, 1:15k). Quadrant 4 did not receive any compound. The following day, 50 µl of fresh medium was added to each well. On day 7 (iT day 21), the media volume was reduced to 50 µl per well using a BioTek plate washer (travel rate 7, height 63) followed by addition of 10 µl 6x staining cocktail containing Calcein Violet AM (Thermo Fisher; final dilution: 1:2,000 = 500nM), anti CD56-FITC (Biolegend; clone 5.1H11; 1:500), anti-CD7-PE (Becton Dickinson; clone M-T701; 1:500), anti-CD4-APV (Biolegend; clone OKT4; 1:200) and 7AAD (BD Biosciences 51-68981E; 1:100) in PBS-/-, mixing with an Apricot Personal Pipettor (384w 125 µl tips, 10 mix cycles, mix volume of 30µl) and incubation for 30 min at room temperature. The staining solution was diluted by removing 60 µl per well using a BioTek plate washer, adding 50 µl of PBS, and removal of 50 µl, with brief plate spin steps (30s at 300xg) performed prior to each plate washer aspiration step. The stained plate was imaged on a Yokogawa CellVoyager 7000 imager (10x lens, 2x binning, confocal mode, 2×2 fields per well, each laser (when used) at 100% power; C01 (Calcein) = 405nm excitation, BP445/45, 1000ms; C02 (FITC) = 488nm, BP525/20, 1000ms; C03 (PE) = 561nm, BP580/23, 1000ms; C04 (APC) = 640nm, BP661/20nm, 2000ms; C05 (7-AAD) = 561nm, BP661/20nm, 1000ms).

Image processing (including channel alignment and background subtraction) and ROI identification and measurements were performed with ImageJ/FIJI (Schindelin et al., 2012) using script iT_V4 with slice median BG subtraction.ijm. To compensate for PE signal bleed-through into channel C05, channel C06 (=7-AAD compensated) was calculated by subtracting 10% of the C02 signal from the C05 signal. Measurement results were analyzed using Python in Google Colab. For screening validation experiments drugs were either added once at the iT day 14 ProT cell stage at the following concentrations (GNF351 1 µM, CH223191 1 µM, EPZ004777 1 µM, OAC1 5 µM, CHIR99021 3 µM) or replenished with every half media change (every 3-4 days).

### Flow cytometry

For *in vitro* differentiation experiments, 50-100 µl of sample were collected from wells and filtered through a filter-top polystyrene flow cytometry tube, pelleted at 300g for 5min and resuspended in flow cytometry buffer (PBS+ 2%FBS) containing the desired antibody cocktail (antibodies are listed in table S2). After 20 min incubation at room temperature in the dark, cells were washed with flow buffer and resuspended in flow buffer containing viability dye (PI, DAPI or Sytox Blue). Sample acquisition was performed on a SONY MA900, LSRFortessa (BD) or Cytek Aurora and analyzed using FlowJo v10.10.0. For zebrafish experiments, pools of zebrafish embryos/larvae were dechorionated and resuspended in 500 ml Liberase TM (Millipore Sigma) solution (75 mg/ml in 1xPBS/1mM EDTA), incubated at 34C and dissociated with a 1000 ml pipette every 15-30 minutes until embryo trunks are no longer visible (Frame et al., 2020); each data point represents 6 zebrafish. CD41^+^ zebrafish HSPCs were gated separately from the CD41hi thrombocyte population (Lin et al., 2005; Ma et al., 2011).

**Table S2.**

| Target | Fluorochrome | Vendor | Cat. No. | Clone |
| --- | --- | --- | --- | --- |
| CD3 | PE-Cy7 | Biolegend | 344816 | SK7 |
| CD4 | PE-Cy5 | BD | 555348 | RPA-T4 |
| CD5 | BV510 | BD | 563381 | UCHT2 |
| CD7 | PE | BD | 555361 | M-T701 |
| CD8 | BV421 | BD | 562428 | RPA-T8 |
| CD45 | APC-Cy7 | BD<br>Biolegend | 557833<br>304014 | 2D1<br>HI30 |
| TCRa/$\beta$ | APC | Biolegend | 306717 | IP26 |
| CD56 | FITC | Biolegend | 362546 | 5.1H11 |
| CD19 | PE | Biolegend | 302207 | HIB19 |

### B cell differentiation

To facilitate B cell differentiation, MS5-hDLL1 stromal cells were seeded onto gelatin-coated 24-well plates and allowed to grow to confluence in MyeloCULT™ H5100 (STEMCELL Technologies #5150) with Penicillin-Streptomycin (Corning #30-002-CI). Once confluent, wells were seeded with 5000-8000 human cord blood CD34^+^ cells in duplicate and cultured in MyeloCULT™ H5100 containing 50ng/mL human recombinant stem cell factor (STEMCELL Technologies, #78155.1), thrombopoietin STEMCELL Technologies, #78210.1), interleukin 7 (STEMCELL Technologies, #78053), and 10 µg/ml FMS-like tyrosine kinase 3 ligand STEMCELL Technologies, #78009) for 4 weeks with media changes twice weekly. AHR antagonists GNF351 and CH223191 were each delivered to cells at a final concentration of 1 µM, and AHR agonist FICZ was delivered at 0.2 µM. DMSO vehicle was included as a control at a final concentration of 1:2500 in culture medium. All compounds were refreshed daily. After 4 weeks, the cells were removed from the wells by incubation with trypsin (Thermo Fisher, #25200056) at 37°C for ten minutes. The cells were then filtered through a filter-top polystyrene flow cytometry tube (Fisher Scientific, #08-771-23), washed with flow cytometry buffer (PBS + 2% FBS + antibiotics), and centrifuged at 300xg for 5 minutes followed by antibody staining and flow cytometric analysis for expression of CD45 and CD19.

### BiTE experiments

All BiTE experiments were performed in 96 U-bottom well plates as follows. Luciferase-expressing target cells were pre-plated at 10k per well in 50 µl of R10. Mature iT cells were added at indicated E:T ratios in 50 µl of R10, followed by BiTE addition. Cells were co-cultured for 48-72 h after which they were lysed by addition of 100 µl lysis and substrate buffer from the Bright-Glo Luciferase Assay system (Promega) and readings were performed on a SynergyNeo (BioTek) plate reader.

Mature iT cells were harvested and replated in R10+IL-7, IL-2 and IL-15 3 days prior to setting up the BiTE experiment, to adjust cells to culture in serum containing medium. Where indicated, effector cells were pre-sorted for live, CD8^+^CD56^-^ to remove unspecific lysis from off-target cells present in the cultures.

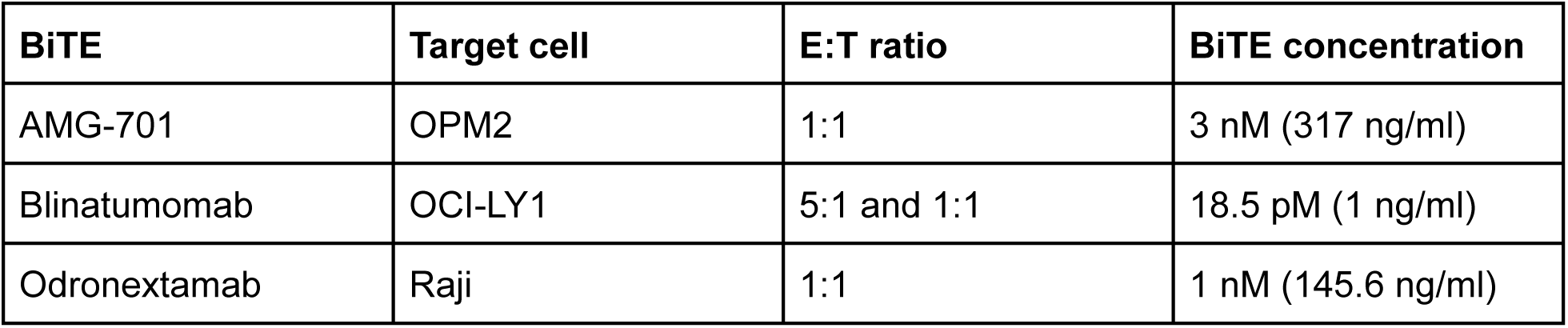

| BiTE | Target cell | E:T ratio | BiTE concentration |
| --- | --- | --- | --- |
| AMG-701 | OPM2 | 1:1 | 3 nM (317 ng/ml) |
| Blinatumomab | OCI-LY1 | 5:1 and 1:1 | 18.5 pM (1 ng/ml) |
| Odronextamab | Raji | 1:1 | 1 nM (145.6 ng/ml) |

### Bulk RNA Sequencing

Total RNA was isolated from CB CD34^+^ derived iT cells on iT day 15 treated with small molecules or DMSO for 24 h (Direct-zol MicroPrep, ZYMO). Differential gene expression analysis and GO term enrichment were performed using NovoMagic, a free Novogene platform for data analysis.

### Zebrafish

Zebrafish maintenance and experiments were performed according to guidelines set forth by the Beth Israel Deaconess Medical Center and Boston Children’s Hospital Institutional Animal Care and Use Committees. Adult zebrafish were housed in a standard circulating water system at 27 °C and fed a brine shrimp diet. AB strains were used for wild-type studies and outcrosses. Transgenics were identified by fluorescence microscopy. For embryonic studies, males and females were separated off-system overnight, and barriers removed the following morning to allow for timed embryo generation. After collection, embryos were maintained under static conditions in E3 embryo buffer in incubators at 28.2 °C to desired developmental stages reported in figure legends. Embryos exhibiting developmental abnormalities or delay were discarded. Sex is not determined in zebrafish embryos and therefore was not assessed.

### AHR Reporter Fish Line Generation

The Tg(8XRE:gfp) line was generated by insertion of an 8x concatemer of the AHR/ARNT binding site (gene block IDT) into a Gateway first entry vector (p5E), and subsequent multi-site Gateway recombination (Invitrogen) into a destination vector carrying the GFP transgene using the Tol2Kit workflow (plasmids listed in Table S3) (Kwan et al., 2007). Germline transgenesis was achieved in 11 founders via co-injection of the construct with Tol2 transposase mRNA.

P5 Insert parts:

Core site: 5’-GCGTG-3’

Binding site: 5’-T/GNGCGTGA/CG/CA-3’

Chosen site: 5’-TAGCGTGCGA-3’

Spacer sequence: TGTCTCAAGCTTGCAG

Minpromoter: AGAGGGTATATAATGGAAGCTCGACTTCCAG

Full 8x sequence insert sequence: aaaaaggtaccACCTGCAGATCTCTAGAAGCTTGCAGTAGCGTGCGATGTCTCAAGCTTGCAGTAGCGTGCGATGTCTCAAGCTTGCAGTAGCGTGCGATGTCTCAAGCTTGCAGTAGC.GTGCG ATGTCTCAAGCTTGCAGTAGCGTGCGATGTCTCAAGCTTGCAGTAGCGTGCGATGTCTCAAGCTTGCAGTAGCGTGCGATGTCTCAAGCTTGCAGTAGCGTGCGActgcaagcttctagagatctgcaggtcgaggtcgacggtatcgatGGATCCAGAGGGTATATAATGGAAGCTCGACTTCCAGCTCGAccgcgg aaaaa

**Table S3.**

| Recombinant DNA | Source | Identifier |
| --- | --- | --- |
| Plasmid: p5E-8XRE | This paper |  |
| Plasmid: pME-GFP | Kwan et al., 2007 | Tol2kit v1.2 #237 |
| Plasmid: p3E-polyA | Kwan et al., 2007 | Tol2kit v1.2 #302 |
| Plasmid: pDestTol2pA2 | Kwan et al., 2007 | Tol2kit v1.2 #394 |

### Zebrafish Embryo Exposure

Groups of 6 stage-matched embryos were arranged in 12 well plates in 2mL E3 water with or without chemical treatments of interest. Zebrafish were dosed with DMSO (1um), FICZ (10nm), CH223191 (1um) and/or OAC1 (1um).

### Statistics

Statistic tests were performed using GraphPad Prism Version 11.0.0. Statistical significance is indicated as *: P ≤ 0.05, **: P ≤ 0.01, ***: P ≤ 0.001, ****: P ≤ 0.0001. For comparison of two groups unpaired t-tests with Welch’s correction were used, in cases where more than 2 groups were compared Ordinary one-way ANOVA was performed.

## ACKNOWLEDGMENTS

This work was supported by NIH fund NIDDK R01DK134515 (RGR), 1R01HL154580 (TEN), and 1RC2DK120535-01A1 (TEN, GQD). Funding for the Cell Voyager imager was provided by the Massachusetts Life Science Center (MLSC project: Stem Cell Imaging and Therapeutics Center). We thank the Eric Smith lab at Dana-Farber Cancer Institute for kindly sharing AMG-701.

## AUTHOR CONTRIBUTIONS

CK and TMS conceived the project and designed the experiments CK, NK, RW, KY, YZ, SW performed the hiPSC and iT experiments. MTW performed zebrafish experiments. KF performed B cell differentiation experiments. MN and RW helped with experiments. MF helped with RNAseq analysis. CK and TMS analyzed the data and wrote the manuscript. TMS, TEN, GQD, and RGR supervised the research, and provided funding.

## DECLARATION OF INTERESTS

Part of this work has been generated under sponsored research agreement with Elevate Bio (Waltham, MA). CSK, GQD, and TMS are inventors on a patent application filed by Boston Children’s Hospital related to methods to enhance production of lymphoid cells from human pluripotent stem cells. The other authors declare no competing interests.

---

**FIG S1.**
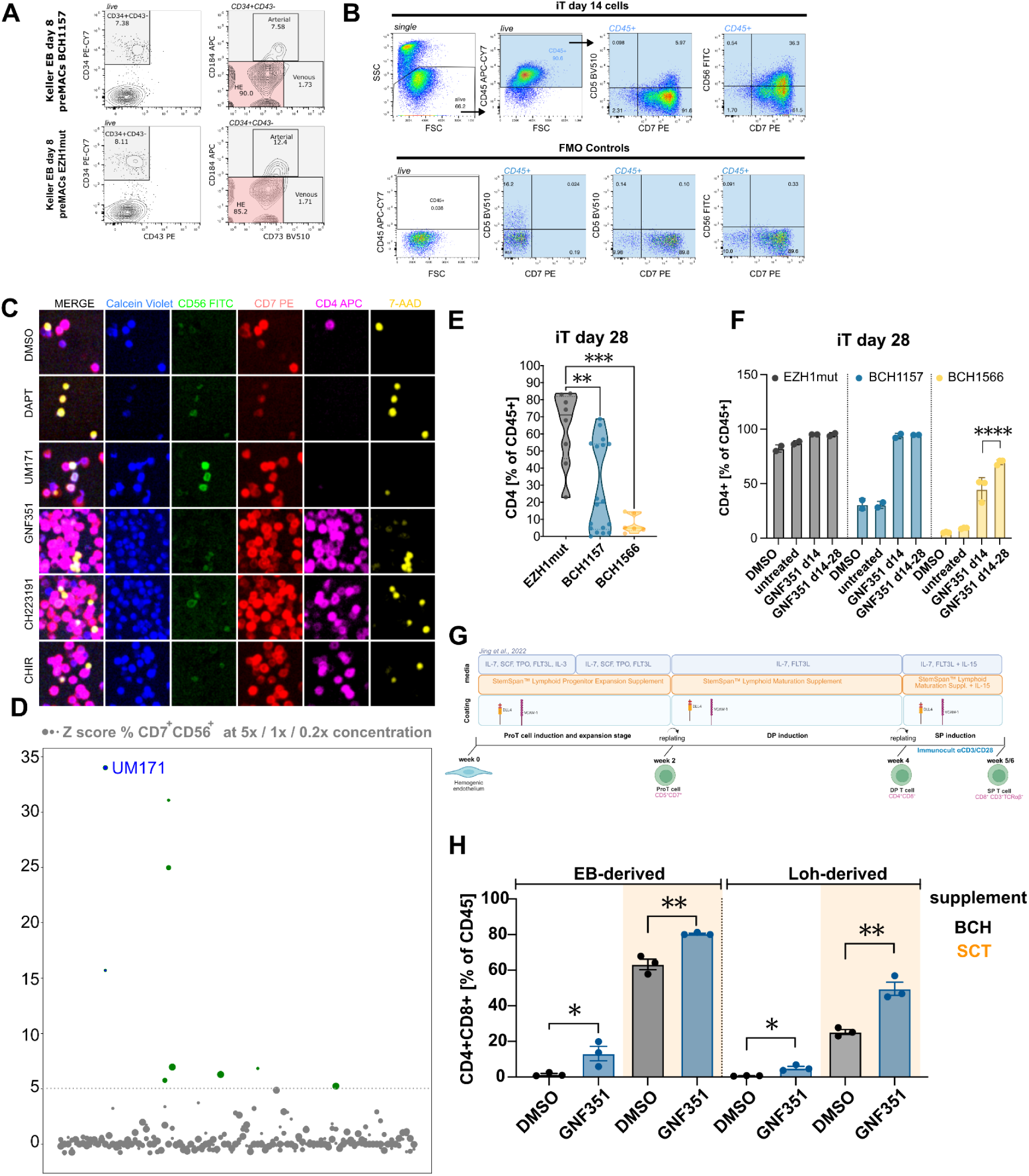
A: Representative flow cytometric profiling of hiPSC-derived hemogenic endothelial cells (CD34+CD43-CD184-CD73-) from BCH1157 and EZH1mut hiPCSs prior to CD34+ MACS enrichment. B: Exemplary gating strategy for flow cytometric analysis of iT day 14 cells. C: Fluorescent-microscopic image of iT day 21 cells after antibody staining against CD56 (FITC), CD7 (PE), CD4 (APC) including viability dyes Calcein Violet and 7-AAD. Scale bar indicates 20 µM. D: Z scores for CD56-inducing hits (potential NK cell inducing agents). Compounds with Z-scores higher than 5 were considered hits. E: Percentage of CD4+ cells on iT day 28 in 3 hiPSC lines. Each data point represents one well of iT cell differentiation. F: CD4^+^ percentage in iT differentiation day 28 cultures from 3 hiPSC treated with GNF351 either as a single dose on iT day 14 or continuously from iT day 14 to 28. Bar graphs depict mean with SEM. G: Schematic of iT differentiation protocol timeline using commercial media vs media published in Jing et al. 2022, H:GNF351 has a positive effect on CD4^+^CD8^+^ independent of HSPC protocol and iT differentiation medium. Percentage of CD4^+^CD8^+^ cells on iT day 28 in hiPSC line BCH1157 using different protocols for the derivation of hematopoietic progenitors and different iT differentiation culture conditions.

**FIG S2.**
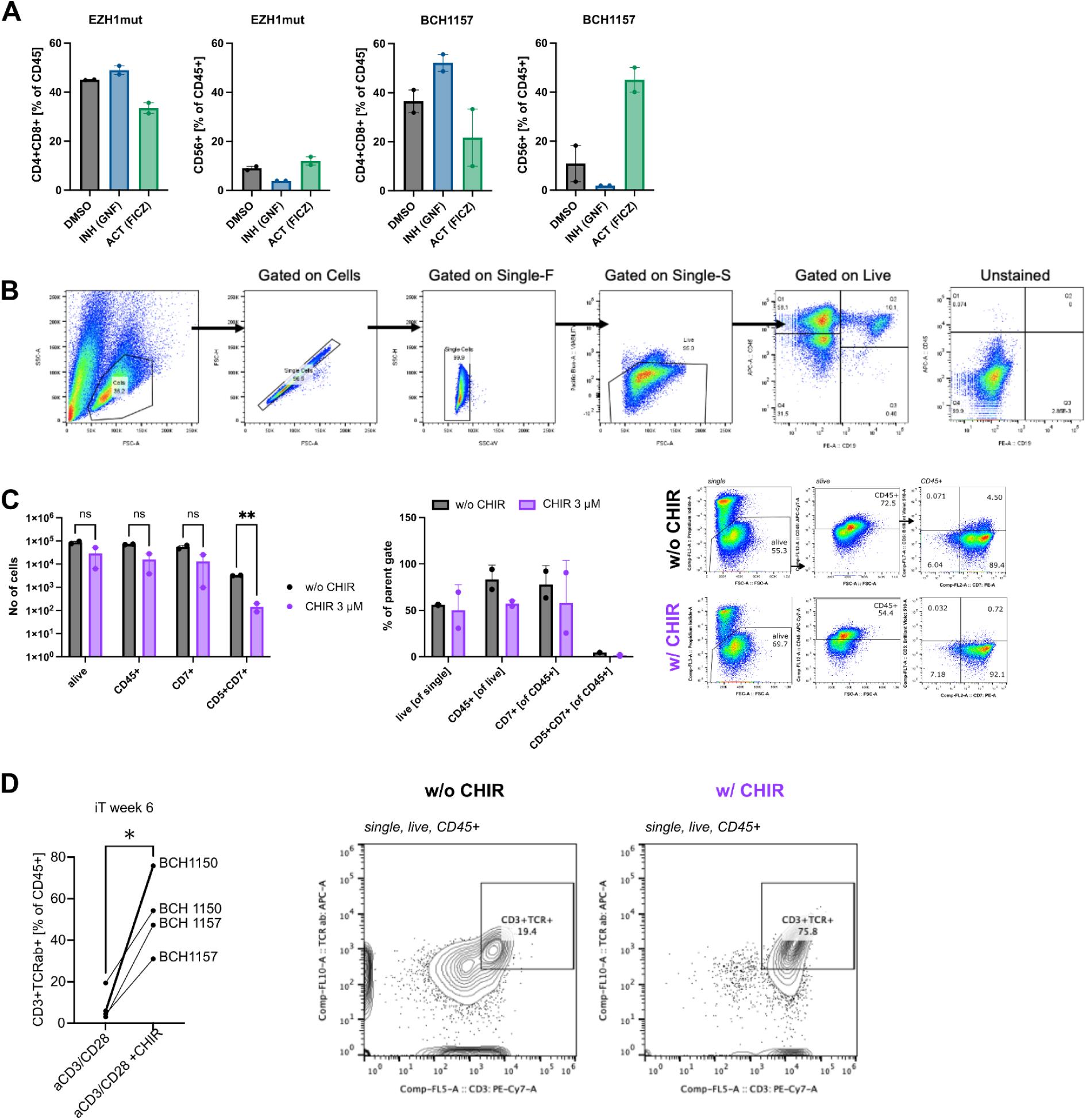
Importance of timing in AHR and WNT modulation. A: Flow cytometric quantification of CD4^+^CD8^+^ or CD56^+^ expression at 4 weeks of iT differentiation in the presence of AHR inhibitor (GNF, 1 µM) or AHR activator (FICZ, 0.2 µM) in two hiPSC lines (EZH1mut or BCH1157). B: Gating strategy for flow cytometric assessment of CB CD34^+^ differentiation outcome towards B cells. C: Flow cytometric profiling of hiPSC to ProT cell differentiation with and without CHIR addition. C: Percentage of CD3^+^TCRab^+^ cells after single positive induction with anti CD3/CD28 antibody with and without CHIR addition.

**FIG S3:**
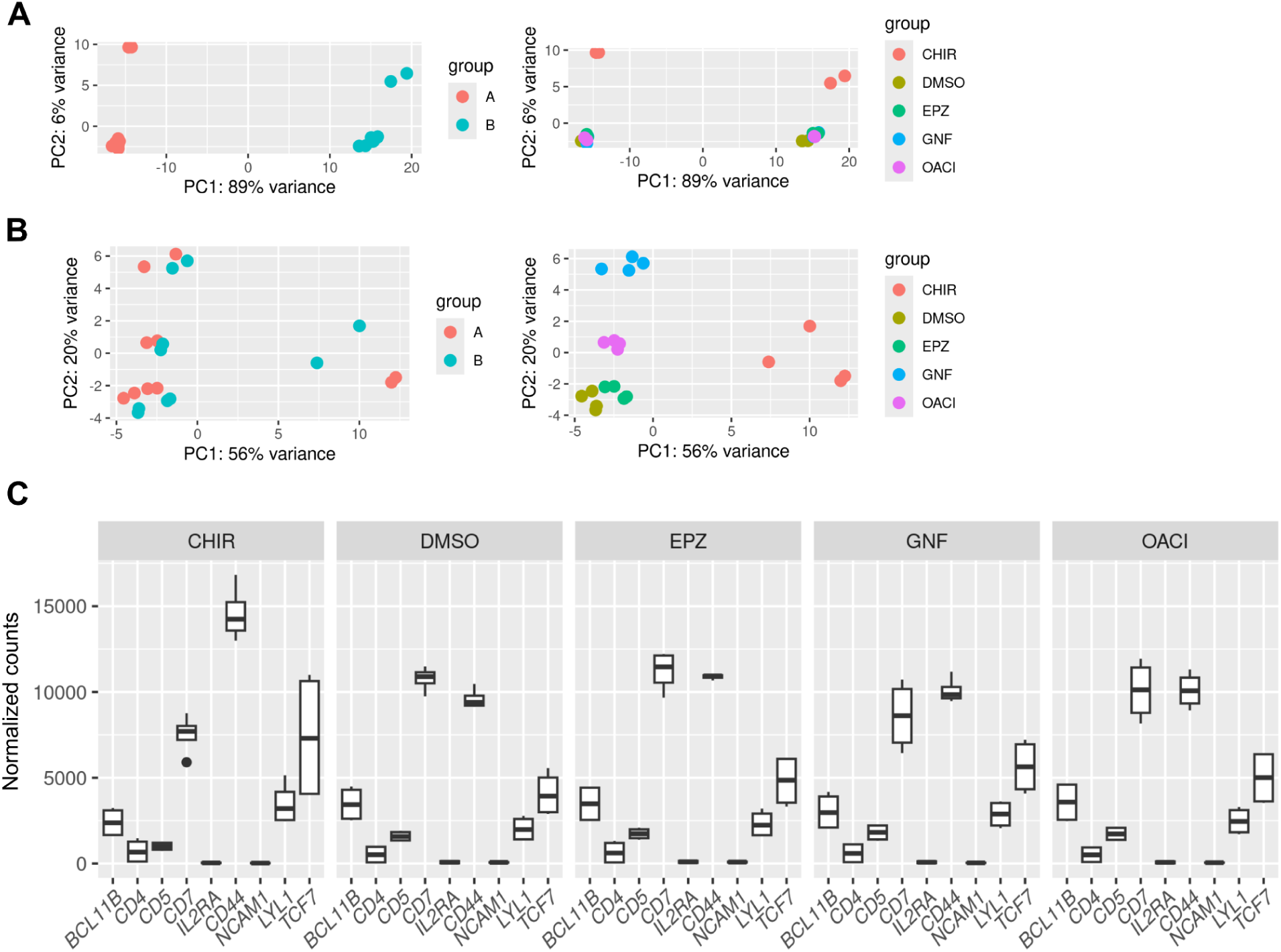
Related to Figure 3. A: Principal component analysis (PCA) of drug treated ProT cells derived from cord blood donor A and B before batch correction and after (B). C: Expression level of ProT cell genes 24h post small molecule treatment

**FIG S4:**
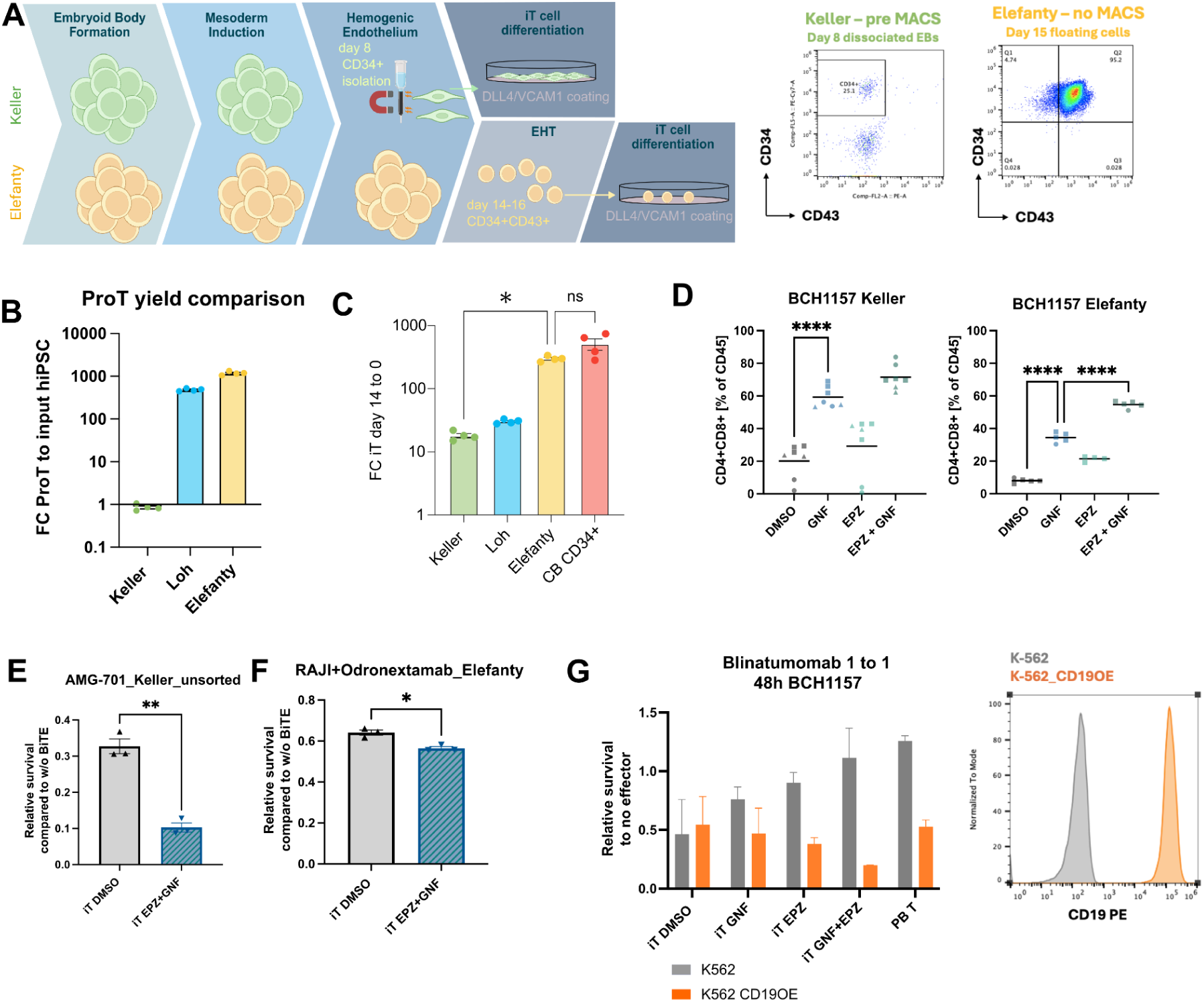
Related to Figure 4. A: Schematic timeline of Keller compared to Elefanty EB differentiation and representative flow plots of input cells for IT differentiation. B: Quantification of fold increase of ProT cells versus input hiPSCs for Keller, Loh and Elefanty hematopoietic differentiation protocols. C: Quantification of fold expansion of plated day 0 cells compared to cell counts on iT day 14 for indicated differentiation protocols and CB CD34^+^ cells. D: Additional experimental data for FIG 4B showing percentage of CD4^+^CD8^+^ double positive cells differentiated from hiPSC line BCH1157 using either the Keller or Elefanty protocol. Each data point represents one well, data from the same differentiation round are indicated by the same shape. One-way ANOVA. E: Relative survival of target cells to no BiTE ctrls after 48h of co-culture with iT cells derived in the presence of small molecule inhibitors without prior enrichment for CD8 SP cells. AMG-701 was added at 3nM F: Relative survival of target cells (Raji) to no BiTE ctrls after 48h of co-culture with iT cells derived in the presence of small molecule inhibitors without prior enrichment for CD8 SP cells. Odronextamab was added at 1nM. G: Relative survival to no effector ctrl of iT cells derived with and without small molecule treatment after 48h of co-culture with K-562 (grey) or K-562 CD19 overexpressing cells (orange) in the presence of 1ng/ml Blinatumomab. Data are from triplicate wells. Flow plot shows CD19 staining of K-562 and K-562 CD19OE cells.

**Supplemental Table S1.**

| agent | stock solvent | vendor | stock concentration [μM] |
| --- | --- | --- | --- |
| 16,16dmPGE2 | DMSO | tocris | 10000 |
| 53AH | DMSO | cellagen | 6000 |
| 740 Y-P | DMSO | medchemexpress | 5000 |
| A-769662 | DMSO | tocris | 50000 |
| AF12198 | DMSO | tocris | 5000 |
| Afatinib | DMSO | selleckchem | 5000 |
| AICAR | DMSO | tocris | 150000 |
| AM580 | DMSO | tocris | 2500 |
| AS101 | DMSO | tocris | 12500 |
| ATRA | DMSO | sigma | 5000 |
| BAI1 | DMSO | tocris | 5000 |
| Bax inh pep V5 | PBS | APEX Bio | 30000 |
| bpV (HOpic) | DMSO | selleckchem | 5000 |
| Calcitriol | DMSO | APEX Bio | 10000 |
| Capsaicin | DMSO | tocris | 5000 |
| CAY10566 | DMSO | caymanchem | 250 |
| Celecoxib | DMSO | selleckchem | 2500 |
| CH223191 | DMSO | selleckchem | 2500 |
| CHIR99021 | DMSO | sigma | 7500 |
| Chroman1 | DMSO | medchemexpress | 250 |
| Cintirorgon | DMSO | medchemexpress | 5000 |
| cyclosporin H | DMSO | sigma | 5000 |
| DAPT | DMSO | APEX Bio | 5000 |
| Dexamethasone | DMSO | tocris | 2500 |
| DMOG | DMSO | selleckchem | 100000 |
| Eltrombopag | DMSO | SCT | 10000 |
| Emricasan | DMSO | medchemexpress | 5000 |
| Eptifibatide | DMSO | sigma | 5000 |
| EPZ004777 | DMSO | selleckchem | 2500 |
| Ferrostatin-1 | DMSO | medchemexpress | 2500 |
| Forskolin | DMSO | tocris | 5000 |
| GDC-0449 (Vismodegib) | DMSO | APEX Bio | 5000 |
| GDC-0941 | DMSO | selleckchem | 1000 |
| GNF351 | DMSO | sigma | 2500 |
| Go 6983 | DMSO | selleckchem | 12500 |
| GSK690693 | DMSO | selleckchem | 500 |
| GW0742 | DMSO | tocris | 425 |
| GW4869 | DMSO | Sigma | 10000 |
| HX 531 | DMSO | caymanchem | 12500 |
| IBMX | DMSO | selleckchem | 50000 |
| IDF-11774 | DMSO | selleckchem | 10000 |
| iMAC2 | DMSO | tocris | 10000 |
| ingenol | DMSO | selleckchem | 25000 |
| ionomycin | DMSO | tocris | 2500 |
| KF 38789 | DMSO | tocris | 15000 |
| L-NAME | DMSO | tocris | 25000 |
| LDN193189 | DMSO | sigma | 6250 |
| Leupeptin hemisulfate | DMSO | tocris | 50000 |
| LY294002 | DMSO | selleckchem | 12500 |
| LY2955303 | DMSO | tocris | 10000 |
| LY364947 | DMSO | selleckchem | 6250 |
| MCC950 | DMSO | selleckchem | 500 |
| Mifepristone | DMSO | tocris | 10000 |
| Nigericin | DMSO | selleckchem | 250 |
| NSC13128 (NAM) | DMSO | selleckchem | 187500 |
| NUTLIN-3A | DMSO | medchemexpress | 10000 |
| OAC1 | DMSO | tocris | 12500 |
| PD169316 | DMSO | selleckchem | 5000 |
| PD173074 | DMSO | selleckchem | 1000 |
| PHA-L | PBS | invitrogen | 500 |
| Pifithrin-α | DMSO | tocris | 10000 |
| Pifithrin-μ | DMSO | tocris | 10000 |
| PKI 14-22 amide,<br>myristoylated | DMSO | tocris | 2500 |
| PMA | DMSO | selleckchem | 2500 |
| PP1 | DMSO | selleckchem | 5000 |
| PP242 | DMSO | selleckchem | 2500 |
| PS48 | DMSO | sigma | 25000 |
| PSB-12379 | DMSO | medchemexpress | 5000 |
| Pyridone 6 | DMSO | medchemexpress | 500 |
| (R)-(+)-Bay-K-8644 | DMSO | medchemexpress | 10000 |
| rapamycin | DMSO | selleckchem | 2500 |
| refametinib | DMSO | selleckchem | 2500 |
| Ro5-3335 | DMSO | tocris | 10000 |
| SAG | DMSO | selleckchem | 250 |
| SANT-1 | DMSO | selleckchem | 1000 |
| SB203580 | DMSO | tocris | 10000 |
| SB431542 | DMSO | tocris | 18000 |
| SB590885 | DMSO | selleckchem | 2500 |
| SC79 | DMSO | selleckchem | 10000 |
| SR 0987 | DMSO | selleckchem | 12500 |
| SR 7826 | DMSO | tocris | 2500 |
| ST2825 | DMSO | medchemexpress | 20000 |
| T0070907 | DMSO | tocris | 2500 |
| tat-BECN1 | DMSO | selleckchem | 12500 |
| trans-ISRIB | DMSO | tocris | 1750 |
| Tranylcypromine | DMSO | tocris | 5000 |
| TRULI (LATS-Inh1) | DMSO | selleckchem | 10000 |
| UM171 | DMSO | selleckchem | 400 |
| UNC0224 | DMSO | tocris | 5000 |
| Verteporfin | DMSO | selleckchem | 12500 |
| Y27632 | DMSO | tocris | 5000 |
| YODA1 | DMSO | selleckchem | 2500 |
| ZK159222 | DMSO | caymanchem | 1000 |
| ZK164015 | DMSO | tocris | 1000 |
| β-Estradiol | DMSO | sigma | 1250 |

